# Distinct vocal flexibility encodes food identity in marmoset monkeys

**DOI:** 10.64898/2026.04.24.720623

**Authors:** Elena Cavani, Alice Chevrollier Oriá, Steffen R. Hage

## Abstract

Combinatorial vocal behavior, in which discrete vocal elements are recombined to expand the vocal repertoire, is a hallmark of human language and has been observed in some nonhuman primates. Here, we show that marmoset monkeys produce distinct vocalizations that discriminate food categories and items by producing long sequences dominated by a single call type whose spectral and temporal features vary. These sequences carried reliable information about food type that could be accurately predicted by trained classifiers. Moreover, rather than combining distinct call types, marmosets rely on sequencing and graded acoustic variation within a single call class to generate functionally diverse signals. These results suggest that precursors of linguistic combinatoriality and multiparameter coding are not unique to human speech but are present in nonhuman primate communication.

## Introduction

Vocal communication enables social animals to coordinate, compete, and cooperate, yet most species achieve this richness of function with a strikingly limited set of vocal elements (Cheney and Seyfarth, 2018; Xie et al., 2024). Since the spontaneous invention of new call types is rare in most species, many vocal systems expand their range by reusing and recombining elements from a limited vocal repertoire (Ouattara et al., 2009; Engesser and Townsend, 2019). Human language appears to have evolved under similar pressures (Hage, 2024). Language is built from discrete meaningless vocal units, such as phonemes and syllables, that are recombined to produce an infinite set of meaningful vocal elements, such as words and sentences (Pleyer et al., 2025). Thus, a finite set of syllables can give rise, through recombination, to an effectively unbounded number of words and meanings(De Boer et al., 2012).

This combinatorial property of vocal communication is rare in non-human primates, but has recently been observed in great apes, and a few Old-World monkey species. Studies show that chimpanzees (Leroux et al., 2023) and bonobos (Berthet et al., 2025) engage in combinatorial vocal behavior, using different vocal elements in a syntax-like manner to signal distinct meanings. In chimpanzees, ordered vocal sequences with combinatorial properties occur at high rates (Girard-Buttoz et al., 2022), and are deployed in ecologically relevant contexts such as predator encounters (Leroux et al., 2023), and Campbell’s monkeys exhibit context-specific combinatorial signaling by arranging vocalizations into structured sequences (Ouattara et al., 2009), further supporting the view that recombination of vocal elements is not uniquely human. These findings rekindle a long-standing debate over whether such language-like capacities evolved only within the hominid lineage or were already present earlier in the primate lineage (Jarvis, 2019; Hage, 2024). Together, they suggest that at least some linguistic principles may be present in non-human primates and could emerge through modulation of preexisting vocal structures.

Marmoset monkeys provide an important test case for this hypothesis. This highly vocal New World monkey species, with a rich vocal repertoire (Martins Bezerra et al., 2008; Agamaite et al., 2015) and flexible use (Pomberger et al., 2019), has recently been shown to exhibit patterns of vocal usage previously considered unique to human language. For instance, marmosets adopt vocal compression strategies such that commonly used vocalizations become abbreviated, consistent with Zipf law of brevity (Risueno-Segovia et al., 2023). Moreover, marmosets form structured call combinations in many contexts, including social interaction, mobbing, and food-related situations. (Bosshard et al., 2024), yet how they encode contextual information, such as the type of threat or available food, remains unclear. This raises a key question - how do marmosets convey biologically relevant information within the constraints of a limited vocal repertoire?

There are two potential mechanisms for information encoding in marmoset vocalizations: recombining different call types and modulating acoustic features, such as frequency or duration, within a discrete call type. To test how these mechanisms contribute to contextual encoding, we investigated vocal sequences during food presentation, a context in which marmosets produce extended sequences of chirp calls (Vitale et al., 2003). Specifically, we examined how different foods are labeled through the combinations and modulation of calls within these vocal sequences. Our findings reveal that marmosets exhibit a distinct form of combinatoriality, in which sequences are constructed from graded acoustic variation of the single call type, the chirp call. This provides a compelling example of how the vocal repertoire can be extended not by increasing call diversity, but by flexibly reusing and recombining variations within existing calls. Together, these results suggest that combinatorial structure can emerge from within-call acoustic variability, rather than exclusively from combinations of discrete call types.

## Material and methods

### Experimental Animals

For this study we used six common marmoset monkeys (*Callithrix jacchus*), three female and three males, aged 7-12 years and housed at the University of Tübingen, Germany. Animals were usually kept in mixed-sex pairs and were all born in captivity; monkeys N and G, and monkeys D and Q, were each housed together as pairs. The animal room was maintained at approximately 26 C, with humidity between 40 % and 60%, and a 12h:12h light-dark cycle. All monkeys had *ad libitum* access to water and food. The diet consisted of commercial pellets, fruits, vegetables, mealworms, and locusts. All animal handling and experimental procedures were approved by the local authorities (*Regierungspräsidium Tübingen*) and in accordance with the guidelines of the European Community for the care and use of laboratory animals.

### Vocal recordings

All recordings were performed in the animal room within the monkeys’ home cages. When animals were housed in pairs, the partner was temporarily removed from the cage for the duration of the recording. Vocalizations were recorded with a condenser microphone (ME66 with preamplifier K6P, Sennheiser electronic, Wedemark, Germany), which was positioned approximately 10 cm in front of the animal’s cage, and digitized at a sampling frequency of 44.1 kHz using an audio interface (UR22mkII, Steinberg, Hamburg, Germany) connected to a computer running recording software (Audition, Adobe, San Jose, CA, USA). Additionally, video recordings were acquired with a 4K camcorder (HC-VX989, Panasonic Corporation, Japan; sampling frequency, 30 frames per second) to provide a visual record of the sessions.

#### Single food presentation

Each recording session involved presenting a single food item to one focal animal. On each experimental day, two different food items and one control condition were tested in pseudo-randomized order across sessions and days. Test food items were banana (6 g), grapes (6 g), mealworms (10 individuals), locust (1 individual), and egg (6 g). Control conditions consisted of standard pellets (20 g), mixed vegetables (20g; 10 g paprika, 10 g zucchini), and an empty food bowl. For each food type we obtained an adequate sample of vocalizations within 4-6 recordings sessions per condition, providing a total of 132 recordings for subsequent analysis (Monkey D: 16 recordings, Monkey E: 21, Monkey R: 21, Monkey G: 23, Monkey N: 31, Monkey Q: 20).

#### Food Preferences Assessment

To assess each monkey’s food preferences, we conducted a multiple-item, tournament-style choice test (Fig 1A). For each animal, we first identified the preferred item within each category (fruit, insect) and then pitted these preferred items against each other to rank first, second, and third overall choices. Each pairwise comparison was repeated six times, and the positions of food items were varied on every presentation to avoid positional bias. The tournament was run once per day. Call rate, defined as the number of vocalizations produced in the first two minutes after food presentation, was calculated from the single food presentations described above.

**Fig. 1.**
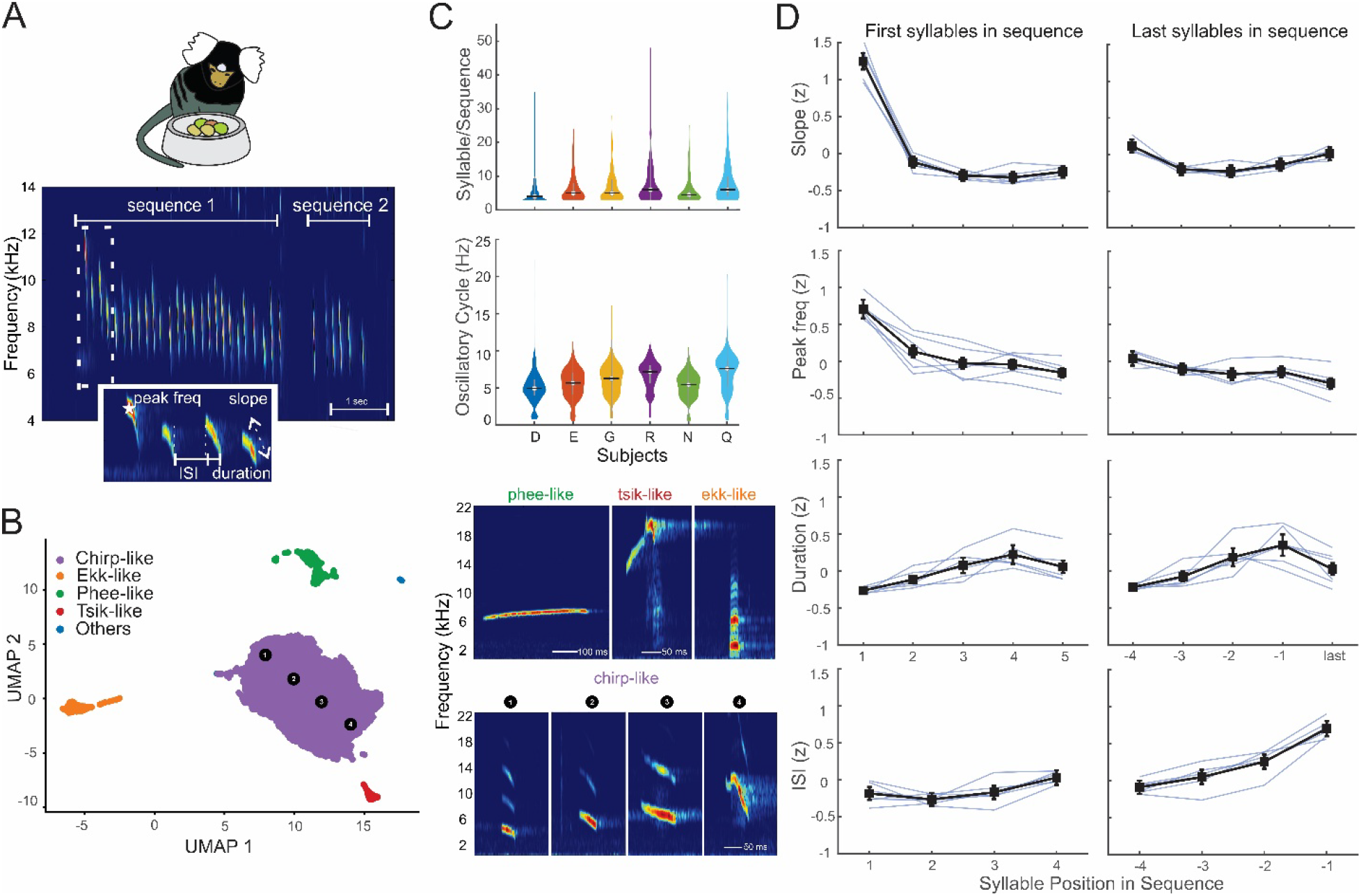
Food-associated calls comprise distinct call types and form vocal sequences with stable structure across animals. **(A)**. Marmoset producing food-associated calls. The spectrogram shows two representative vocal sequences. The inset illustrates extracted acoustic parameters (peak frequency, inter-syllable interval (ISI), duration, and frequency slope). **(B)**. UMAP projections and HDBSCAN clustering of all calls produced during food presentation. Representative spectrograms for each cluster are shown on the right. For the large chirp-like cluster, four representative examples are shown sampled across the cluster **(C)**. Top, violin plots showing the distribution of syllables per sequence for each monkey. Black horizontal bars indicate medians. Bottom, violin plots show the distribution of oscillatory rates across animals. **(D)**. Sequence structure, showing how selected acoustic features (frequency slope, peak frequency, duration, and ISI) change over the course of sequences. Left, first five syllables; right, last five syllables. Black lines indicate mean ± SEM across all animals, single light blue lines indicate individual subjects.

### Acoustic Analysis

We analyzed 9,997 vocalizations recorded in 132 sessions of six marmoset monkeys (Monkey D: 2613 calls, Monkey E: 1085, Monkey G: 1,315, Monkey R: 2,015, Monkey N: 1,524, Monkey Q: 1,445). Call onsets and offsets were manually annotated, and confounding call echoes were carefully identified and removed using standard software (Avisoft-SASLab Pro, Germany). Call duration was defined as the time between vocalization onset and offset. Inter-syllable interval was defined as the time between the end of one vocalization and the onset of the next. The signal was high-pass filtered at 0.5 kHz to remove low frequency disturbances. Spectrograms were computed using a 1024-point FFT with a Hamming window (512 samples) and 50% sample overlap, yielding a frequency resolution of 43 Hz and a temporal resolution of 0.36 ms. For the global call analyses, all vocalizations were included, whereas the sequence analyses were restricted to vocalizations belonging to sequences of at least three vocalizations (8,307 calls).

### Statistical Analysis

All analyses were performed using custom code written in Python and MATLAB. To account for repeated measurements within subjects and to control for subject-related effects, we used linear mixed-effect models (LMMs) to assess relationships between acoustic parameters, sequence structure, and condition. First, we tested for systematic changes in acoustic parameters over syllable position in the sequence (Fig 1D) with the model: Acoustic Feature ∼ Syllable Position + (Syllable Position | Subject). Second, we evaluated overall acoustic features across food conditions (Fig 2D) using: Acoustic Feature ∼ Condition + (Condition | Subject). Finally, we tested for interaction between syllable position and food condition (Fig 2E) with Acoustic Feature ∼ Condition * Syllable Position + (1 | Subject). In all performed tests, significance was tested at an α level of 0.05.

**Fig. 2.**
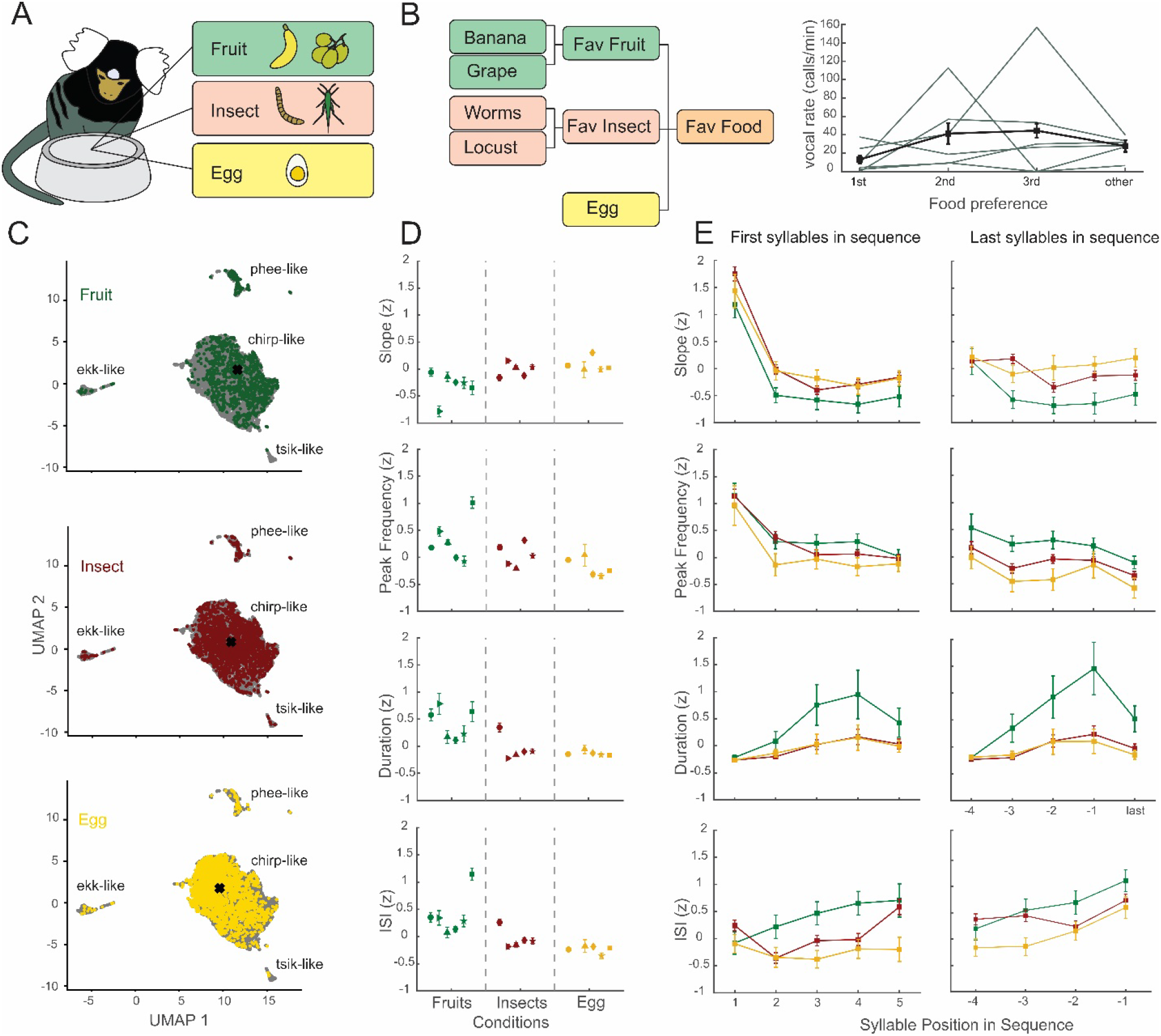
Food-associated vocalizations differ according to food category. **(A)**. Schematic of food conditions presented to each subject: fruit (banana or grapes), insect (mealworms or locust), and egg. **(B)**. Food preference assessment in a tournament-style design (left). Vocal rate (calls/min) during the first 2 min after food presentation is shown for first, second, third preferences, and for “other” (items not chosen as favorites) for each marmoset. Black line and bars indicate mean ± SEM across animals **(C)**. UMAP projections of calls produced under each condition. Black crosses indicate cluster centroids. **(D)** Subject-level means of frequency slope, peak frequency, duration, and ISI. Symbols represent individual subjects (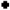 monkey D, 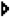 monkey E, 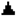 monkey G, 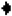 monkey R, 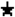 monkey N, 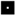 monkey Q), error bars indicate within-subject SEM. **(E)** Sequence structure, showing how acoustic features (slope, peak frequency, duration, ISI) evolve over the course of sequences for each food category. Each line represents, for the given acoustic feature, mean across animals and conditions (green: fruit, red: insect, yellow: egg), bars symbolize SEM.

#### UMAP and Random Forest Classifier analysis

We used Uniform Manifold Approximation and Projection (UMAP)(McInnes et al., 2018) for dimensionality reduction to determine the variety of vocalizations in our data set. Clusters were obtained by applying HDBSCAN (minimum cluster size = 100) to the UMAP embedding, yielding five main clusters. To decode food items and categories (Fig. 3A-C), we trained supervised random forest classifier separately for each subject. Food items with less than 100 vocalizations were excluded from the analysis. To control for class imbalance, we performed random undersampling, so that all classes contained the same number of vocalizations, matching the smallest class. For each iteration, data were randomly split into training (80%) and test (20%) sets. We trained 100 models with a random forest classifier (150 trees, minimum leaf size = 5) and evaluated performance as classification accuracy on the test set. Chance-level performance was estimated with a condition permutation test, in which condition labels in the training sets were shuffled prior model fitting and accuracy was calculated on the held-out test set. This procedure was repeated for 100 random splits, and performance metrics were averaged across iterations.

**Fig. 3.**
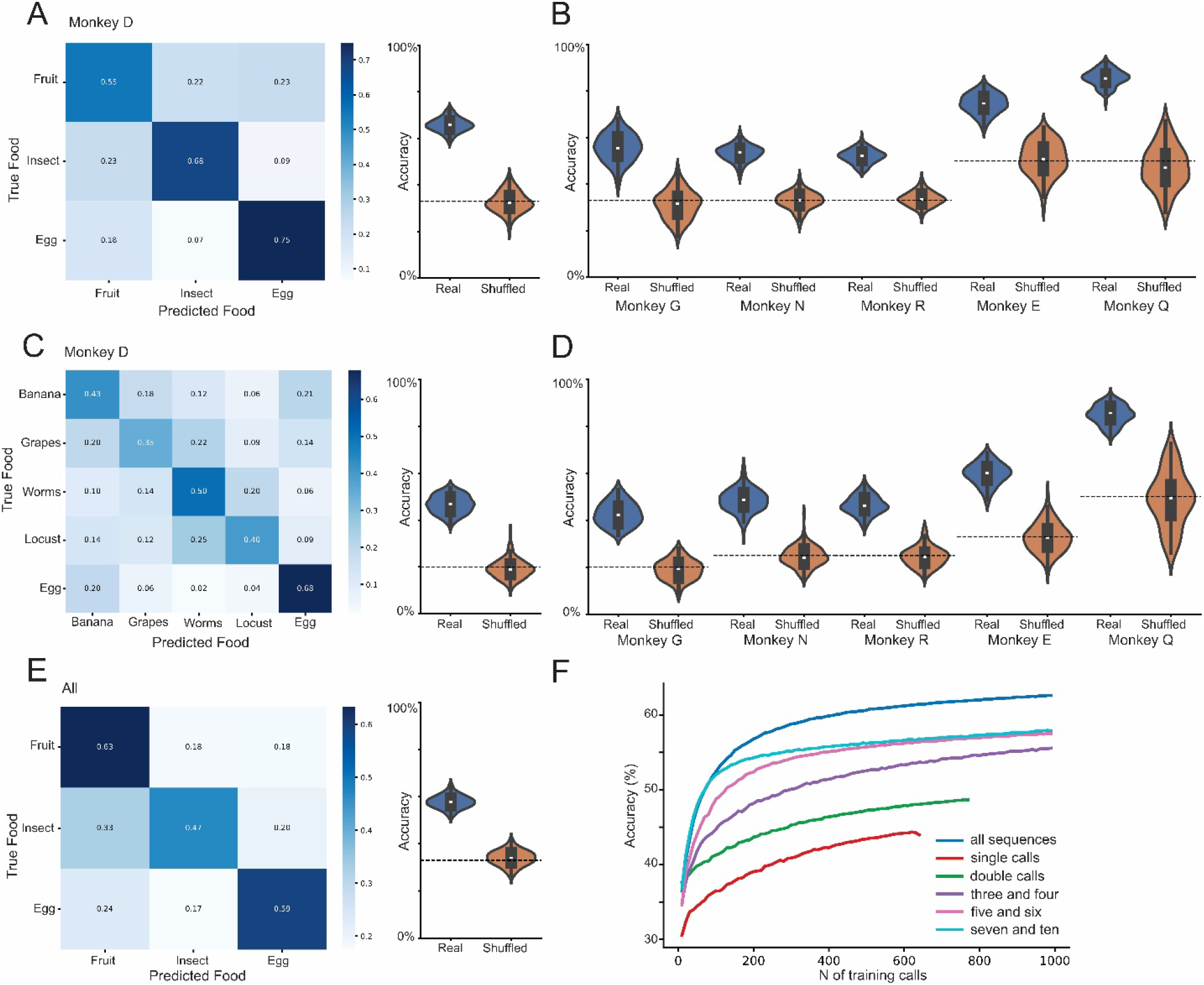
Food-associated vocalizations predict food identity. Confusion matrices of classification accuracy of a 100-model random forest classifier. **(A)** Decoding food categories (fruit, insect, egg) at the single-animal level. For each monkey, the classifier was trained and tested on the individual’s data using 100 bootstrap iterations. Left: Example confusion matrix for monkey D (color-coded classification accuracy). Right: Violin plot showing the distribution of classification accuracies across 100 models for real vs. shuffled data. Horizontal black line indicates chance level. **(B)** Violin plots showing classification accuracies across 100 models for real vs. shuffled data for food category decoding in the other 5 individuals. Horizontal black lines indicate chance. Chance levels differ across animals due to variation in the number of food categories for which they produced vocalizations (3 categories: 33%, 2 categories: 50%). **(C)** Single-animal decoding of individual food items (banana, grape, mealworm, locust, egg). For each monkey, the classifier was trained and tested on the individual’s data using 100 bootstrap iterations. Left: Example confusion matrix for monkey D (color-coded classification accuracy). Right: Violin plot showing classification accuracies across 100 models for real vs. shuffled data. Horizontal black line indicates chance. **(D)** Violin plots showing classification accuracies across 100 models for real vs. shuffled data for single food item decoding in the other 5 individuals. Horizontal black lines indicate chance, which varies with the number of available food categories. Chance levels differ across animals due to variation in the number of food items for which they produced vocalizations (5 items: 20%, 4 items: 25%, 3 items: 33%, 2 items: 50%). **(E)** Population-level decoding of food categories across subjects. For each iteration, 100 vocalizations per subject were sampled, and classification accuracy was computed over 100 bootstraps. **(F)** Sequence-length decoding analysis. Classifiers were trained on vocalization subsets with sequences of different syllable lengths and tested on the full dataset. Colors indicate accuracy for each sequence-length subset.

To predict food items from pooled subjects (Fig 3D), we again used a random forest classifier. Acoustic features were z-scored separately for each subject prior to pooling. For each iteration, we constructed a balanced dataset by randomly sampling 100 vocalizations per condition per subject; only subject-condition combinations with at least 100 vocalizations were included. The sampled data were then pooled across subjects and split into training (80%) and test (20%) sets. As above, we trained a random forest classifier with 150 trees and minimum leaf size of 5. and estimated chance performance with the same permutation procedure.

For the sequence-length decoding analysis (Fig 3E), we adapted an approach previously used for single-neuron data (Elston and Wallis, 2025). We used the same classifier as described above: acoustic features were z-scored per subject, and vocalizations were grouped by sequence length into the following categories: all sequences, single calls, double calls, sequences with 3-4 syllables, 5-6 syllables, and 6-10 syllables. For each iteration, the dataset was split into training (80%) and test (20%) sets using stratified sampling to preserve class proportions. Classifiers were trained on subsets of the training data belonging to each sequence-length category. To assess how decoding performance depended on training data size, we varied the number of training vocalizations from 10 to 1000 in steps of 10, randomly sampling from the corresponding sequence-length category at each step. Classification accuracy was always evaluated on the same fixed test set containing all vocalizations in the dataset. For the UMAP/HDBSCAN and all random forest analyses, we used all vocalizations in our dataset and 19 acoustic features (Table S4).

#### Quantitative analyses of call sequence structure

We calculated Zipf values for each condition, following the approach described previously (Gultekin et al., 2021), to quantify differences in call-type usage across conditions. Briefly, we quantified inter-call and inter-subtype transition probabilities and, for each condition, computed mean transition matrices across subjects to capture how likely one call type or subtype was to follow another. We then summarized overall repertoire organization by deriving a Zipf value from the rank–frequency distribution of call types as a measure of sequence complexity. Zipf values were compared across conditions using a linear mixed-effects model with condition as a fixed effect and subject as a random effect. In addition, we performed chi square tests to detect if the transitions were different between call types.

## Results

We analyzed marmoset vocalizations produced in the presence of several food items: two types of insects, two types of fruit, and egg. We recorded 9,997 food-associated calls from six common marmoset monkeys (*Callithrix jacchus*) in an experimental setting. When presented with food, marmosets produced a high number of calls that were predominantly uttered in sequential bouts (Fig 1A). For each call, we extracted acoustic features including peak frequency, inter-syllable interval (ISI), duration, and frequency slope (peak frequency over time; Table S1). To characterize the call types present in the dataset, we applied UMAP dimensionality reduction followed by HDBSCAN clustering, which revealed five distinct clusters (Fig. 1B). Based on previous quantitative analyses of marmoset call types (Agamaite et al., 2015), we classified these clusters as chirp-like (8751 calls; call percentage/monkey= 89.8% ± 4.5%, n=6), phee-like (543 calls, 5,3% ± 0.8%), tsik-like (182 calls; 1.2% ± 1.1%, n=6), ekk-like (478 calls, 3.3% ± 2.7%, n=6), and other calls (43 calls, 0.4% ± 0.2%, n=6). Therefore, consistent with previous studies, marmosets produced predominantly chirp calls during food presentation, with only a limited number of other call types (Martins Bezerra et al., 2008; Agamaite et al., 2015; Rogers et al., 2018). Notably, they called rarely or not at all during control presentations, which consisted of an empty food bowl, pellets, or vegetables (nine calls from two individuals across all control trials; see Material and Methods).

We next quantified the structure of vocal sequences, defined as in Fig S1, by counting the number of syllables per sequence and summarizing central tendency at the subject level. Median sequence length ranged from 4 to 6 syllables across subjects, with an overall median of 5.1 syllables per sequence, suggesting a consistent sequence structure across individuals (Fig. 1C). Based on this observation, subsequent analyses were restricted to vocalizations belonging to sequences with at least 5 syllables, allowing us to probe sequence structure, and because longer sequences provide more informative examples for characterizing sequence organization. We also calculated the oscillatory rate of all vocalizations belonging to these sequences and found median rates between 5 and 8 Hz across subjects (overall median 6.1 Hz), consistent with previous work on marmoset vocal production (Risueno-Segovia and Hage, 2020) and similar to the phonatory rhythmicity described for human speech (Poeppel and Assaneo, 2020).

Further analysis focusing on sequences revealed systematic changes in acoustic trajectories over sequence position (Fig. 1D). Specifically, we investigated the first 5 syllables and the last 5 syllables of a call sequence to quantify how acoustic features evolved from sequence onset to offset (Table S1). Across sequences, most acoustic parameters differed significantly with syllable position at both the beginning and end of the sequence (for an overview of statistical analyses, see Table S3). For example, frequency slope varied across syllables at the start (F(4,3413)=93. 36, p =1.86e-75, LMM), with the first syllable exhibiting a higher slope than subsequent syllables, and also declined at the end of the sequence (F(4, 3410) = 19.29, p =1.04e-15, LMM). Similarly, peak frequency varied with syllable position (F(4, 3413) = 81.75, p = 2.35e-66, LMM), showing a progressive decline over the course of the sequence and reaching its lowest value in the final syllable. By contrast, temporal features showed the opposite pattern, with both duration and ISI increasing towards later syllables (duration at start: F(4, 3413)=9.61, p= 1.0035e-07; ISI at start: F(4, 3239) = 9.06, p = 2.81e-07; Duration at end F(4, 3410) = 12.74, p = 2.70e−10; ISI at end F(3, 2728)=76.16, p= 2.63e-47, LMM). Taken together, the position-dependent changes in slope frequency and timing indicate that marmoset food-associated call sequences are not acoustically uniform, but instead exhibit orderly within-sequence dynamics that are conserved across individuals.

Next, we asked whether call sequence structure is modulated by the types of food presented. We tested three categories: fruit (banana or grape), insect (mealworms or locust), and egg (Fig. 2A; see Material and Methods). To account for individual preferences, we included two items per category to increase the likelihood that each monkey produced vocalizations for at least one item in each category. For each animal, we first assessed food preferences using a tournament-style paradigm (Fig 2B and Material and methods, Table S2), and then computed call rate during presentation of individual food items. This revealed that call rate did not depend on food preference (Fig. 2B; F(1,127)=0.25, p=0.62, LMM).

We then examined whether the calls produced during food presentation differed according to the food categories offered. In UMAP space, calls occupied partially distinct regions with chirp-like calls showing shifts according to food category, suggesting context-dependent variations in this call type (Fig. 2C). To quantify these differences, we analyzed acoustic features at the subject level. For each monkey, we computed mean frequency slope, peak frequency, duration, and ISI for each category (Fig. 2D). Despite individual variations in food preference, subjects exhibited similar acoustic patterns for each category, consistent with a shared encoding of food category in these vocalizations. Linear mixed-effects models confirmed robust category-dependent differences. Frequency slope differed significantly across food categories (F(2, 6353) = 6.02, p= 0.002, LMM), with fruit-associated calls showing lower slopes than insect- and egg-associated calls. Peak frequency also varied (F(2, 6353) = 9.13, p =1e-04, LMM), with egg condition exhibiting lower frequencies. Duration (F(2, 6353) = 12.64, p = 3.32e-06, LMM) and ISI (F(2, 5670) = 21.038, p = 7.88e-10, LMM) likewise differed, with fruit-associated calls displaying longer durations and ISIs than calls associated with other categories.

Finally, we examined how these features evolved over the course of syllable sequences (Fig 2E). For all parameters, syllable-by-syllable trajectories differed by food category. For example, frequency slope showed a stronger decline across syllables in fruit sequences than insect or egg sequences (F(8,3403)=2.45, p=0.012, LMM). Peak frequency diverged between food categories towards the end of sequences (F(8,3403)= 2.18, p= 0.026, LMM), and ISI increased across syllable positions in the fruit condition, indicating longer ISIs as sequences progressed. Overall, fruit sequences showed stronger within-sequence modulation, whereas insect and egg sequences showed flatter trajectories across several acoustic parameters, potentially reflecting greater variability or flexibility in those contexts. Together, these results indicate that food-related vocalizations and their sequences are structurally organized, consistent across individuals, and systematically modulated by food category.

Having established the existence of robust category-dependent differences, we next asked whether food categories could be decoded directly from the vocal signals. To do so, we trained a 100-model random forest classifier based on 19 acoustic features (Table S4). For each animal, separate classifiers were trained and tested on that individual’s data to predict *i)* food category (fruit, insect, egg; FIG 3A and 3B) and *ii)* individual food items (banana, grapes, mealworms, locust, egg; FIG 3C and 3D). In both tasks, classifiers reliably recovered the food type from vocalizations for all monkeys: food categories were decoded with a mean accuracy of 66% ± 3 % (Monkey D, chance=33%; Fig. 3A), and individual food items with a mean accuracy of 47% ± 3 % (Monkey D, chance=20%; Fig. 3B). Misclassification occurred primarily between items within the same category, consistent with shared characteristics among category members. When data were pooled across animals, food categories remained decodable above chance (58% ± 3%, chance: 33%, Fig. 3E), indicating that food-associated calls share a common acoustic code for category amongst individuals (mean accuracy values per subject 44-81%, mean 60% ± 5%. Because not all animals vocalized for all food items (Fig.3B and 3D), we additionally trained classifiers on the subset of monkeys that produced calls for every category (Figure S2). In this subset, accuracies remained high (52% ± 3%, chance: 33%), and classifiers reliably recovered food type from vocalizations (mean accuracy values per subject 44-61%; mean 52 ± 3.5%). Finally, we trained classifiers on subsets of vocalization sequences with different lengths and single and double calls, compared with the full dataset (Fig. 3F). Classification accuracy increased with sequence length, with longer sequences outperforming single calls, double calls, and short sequences, highlighting the functional importance of producing extended vocal sequences.

We next examined how the vocal repertoire itself was deployed across food conditions. Because all monkeys predominantly produced chirp-like (89.8 ± 4.5%,) and, to a lesser extent, phee-like calls (5,3 ± 0.8%) in response to food, with all other call types being rare (4.8 ± 3.9% combined), subsequent analyses focused on these two dominant categories (Fig S3). Phee calls occurred in response to all food categories but were disproportionally produced in response to fruit compared to insect and egg (Fruit vs Insect: p = 0.038; Fruit vs Egg: p < 0.001; Fig 4A). Chirp-like vocalizations exhibited substantial acoustic variability across sequences (Fig 4B), yet formed a single large cluster in UMAP space, prompting a finer subdivision. Using peak frequency - the main contributor to the UMAP embedding - we defined four chirp-like subtypes (C-1 to C-4, from low to high peak frequency; Fig 4C).

**Fig. 4.**
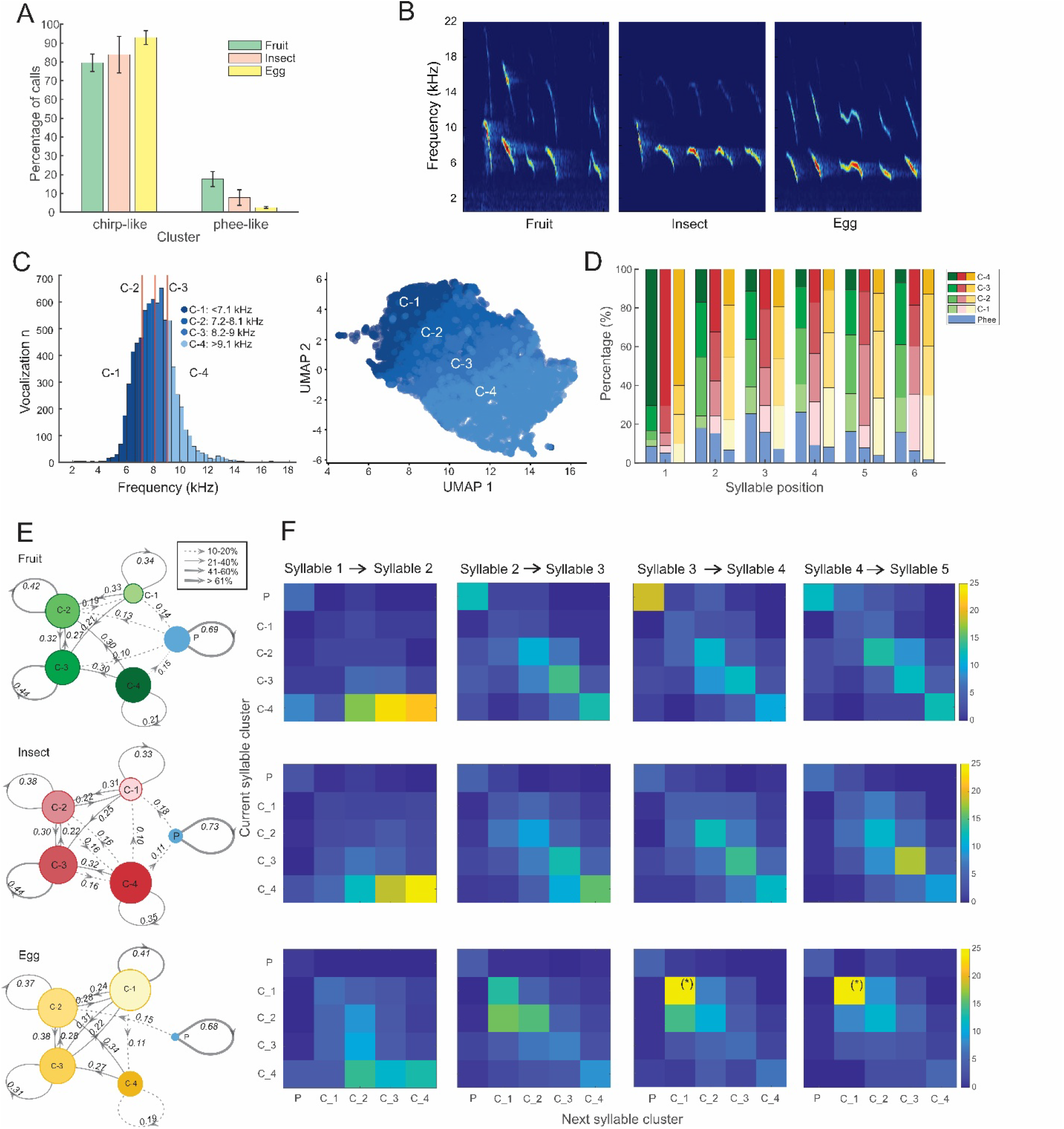
Context-dependent organization and transition structure of marmoset vocal sequences. **(A)** Mean percentage (± SEM) of chirp-like and phee-like calls according to food category across all subjects. **(B)** Example spectrograms of the initial five syllables of vocal sequences produced across the three food categories. **(C)** Subdivision of chirp-like vocalizations into four quartiles based on the maximum peak frequency. Left, histogram illustrates the division into four chirp subtypes (C-1 to C-4, low to high peak frequency). Right, UMAP projection showing the distribution of each chirp-like subtype. **(D)** Stacked bar plot illustrating the mean percentage (across subjects) of phee calls and each chirp-like subtype over the first six syllables of sequences. Phee calls are shown in blue, chirp subtypes for fruit, insect and egg in shades of green, red, and yellow, respectively. **(E)** Transition diagrams for vocal sequences in response to each food category. Nodes represent call types or subtypes, with node size reflecting overall occurrence probability. Arrows indicate transitions, and arrow thickness and style encode transition probability. Transitions probabilities below 10% are not shown for clarity **F)** Transition matrices showing call-to-call transition probability across subjects for syllable transitions 1→ 2, 2→3, 3→4, and 4→5. (*) symbolizes values above 25%.

We then investigated how these chirp-like subtypes were sequentially organized and associated with food condition. In response to each food category, we computed the proportion of each chirp subtype and phee call in the first six syllables of each sequence (Fig 4D). Consistent with our earlier trajectory analyses, higher-frequency chirp variants (C-3 and C-4) were enriched in earlier sequence positions, whereas lower-frequency variants (C-1 and C-2) became more prevalent later in the sequence, especially in response to egg. Phee calls were more likely to occur in later sequence positions. Together, these observations suggest systematic, context-dependent differences in call usage across sequences. We first confirmed a significant effect of condition on the distribution of call types and subtypes (χ^2^(8) = 637.86, p < 0.001); Fig 4D). To probe sequence structure more directly, we quantified inter-call and inter-subtype transition probabilities. For each condition, we computed the mean transition matrices across subjects, capturing the likelihood that a given call type or subtype transitioned to another (Fig. 4E). We further summarized vocal repertoire organization using the Zipf index, a measure for sequence complexity reflecting the rank-frequency distribution of call types. Zipf indices differed significantly across conditions (p = 0.016, LMM; Fig. S4A): fruit-associated sequences showed the highest indices, indicating relatively sparse, even usage of call types and subtypes, whereas egg sequences exhibited the lowest index, consistent with dominance of a smaller subset of calls and reduced repertoire flexibility. To assess differences in sequence structure, we compared transition frequencies across conditions using a chi-square test. This analysis revealed a significant effect of condition on transition structure (χ^2^(48) = 1497.2, p < 0.001, Cramér’s V = 0.335), indicating that transition probabilities varied across contexts. For example, examination of residuals showed that phee to phee and, more generally, phee-associated transitions were overrepresented in the fruit condition. In contrast, C1→C1 transitions were overrepresented in the egg condition, but largely absent in the fruit and insect conditions. Additionally, higher-frequency cluster transitions such as C3→C3, and C4→C4 were underrepresented in the egg condition (Fig S4B and S4C). These effects were further illustrated by examining transition probabilities across early sequence positions (Fig. 4F). As observed earlier (Fig. 2E), initial transitions were dominated by high-frequency chirps, whereas later transitions progressively shifted toward frequency ranges more characteristic of each food category (for example, C1→C1 transitions for egg).

Taken together, these results show that marmosets do not merely modulate acoustic features within a call type, but also flexibly recruit distinct call subtypes and transition patterns to structure vocal sequences that are specific to distinct categories of food. This provides strong evidence that marmosets encode contextual information through both graded acoustic modulation and combinatorial structuring of their vocalizations.

## Discussion

We found that marmosets use their vocal repertoire to reliably differentiate between food categories and items. They did so primarily by assembling extended sequences of a single call type, the chirp-like call, and modulating its acoustic structure in a context-dependent manner, rather than recombining different call types. This shows that food-associated vocalizations in this species are not fixed signals but constitute a flexible system that supports context-dependent variation. Our data provide a compelling example of how marmosets overcome the constraint of a limited vocal repertoire by modulating the acoustic features of existing calls and concatenating them into longer sequences, thereby exhibiting recombinant-like vocal behavior - a feature long regarded as a core design property of human language (Pleyer et al., 2025). Importantly, our results reveal a distinct form of vocal combinatoriality in marmosets. In the food context, marmosets appear to combine variants, or subtypes, of the same call class -sharing core features but differing in specific acoustic parameters – rather than primarily combining different call types. This suggests that combinatorial signaling can operate at multiple levels, even within call-type variation and may therefore more common than previously thought.

What might drive such vocal flexibility? Many non-human primates, including chimpanzees (Slocombe and Zuberbühler, 2006; Déaux et al., 2023), bonobos (Clay and Zuberbühler, 2009), gorillas (Luef et al., 2016), and macaques (Hauser and Marter, 1993) produce food-associated calls that have been linked to properties (such as quantity, quality, and divisibility) or preferences. For instance, increased vocal rates have been linked to food preferences in bonobos (Clay and Zuberbühler, 2009) and chimpanzees (Slocombe and Zuberbühler, 2006). In our data set, however, call rate did not increase for preferred foods, and control presentations with larger quantities elicited few or no vocalizations, arguing against explanations based solely on preference or amount. Instead, the patterns we observe are more consistent with an adaptation to the ecological demands of foraging in small, social prey animals for whom efficient detection, assessment, and sharing of information about food is critical. On this view, vocal flexibility might function to signal the quality and relevance of particular foods within the group.

Moreover, similar acoustic patterns emerged in individuals that were not co-housed and originated from different groups, making it unlikely that our results solely reflect arbitrary, learned labels for specific foods. Rather, they point to a shared mapping between aspects of food value and structured vocal output. Future work will be required to pinpoint which dimensions drive these modulations – for example, whether spectral temporal patterns track nutrient content such as protein or sugar levels. A promising approach will be to introduce novel foods with systematically varied nutritional profiles and ask whether predictable shifts in vocal structure follow.

Our findings also dovetail with work on other marmosets call types. Long-distance phee calls encode identity-related information about intended recipients through coordinated changes along multiple acoustic dimensions (Oren et al., 2024), demonstrating that graded modulation of existing calls can support rich social signaling. Similarly, we find that decoding of food categories depends on the joint contribution of several acoustic features, with frequency-related measures among the strongest predictors. Together, these observations indicate that marmosets routinely exploit multiple degrees of freedom within their vocal repertoire – combining calls into sequences and shaping both spectral and temporal structure – to generate functionally flexible yet interpretable biological signals. This supports the view that core precursors of language, including combinatoriality and multi-parameter acoustic coding, are implemented in non-human primate communication systems.

## Acknowledgments

We thank Avani Koparkar and Thomas Elston for fruitful discussions throughout the preparation of the manuscript.

## Author contributions

Conceptualization: E.C., and S.R.H.; Methodology: E.C., and S.R.H.; Investigation: E.C., and A.C.-O.; Visualization: E.C.; Funding acquisition: S.R.H.; Project administration: S.R.H.; Supervision: S.R.H.; Writing – original draft: E.C., and S.R.H.

## Data and materials availability

All data are available in the manuscript and the supplementary materials or may be obtained from the corresponding author upon reasonable request.

