## Supplemental Figures and Tables for "Distinct vocal flexibility encodes food identity in marmoset monkeys"

### Supplementary Figures

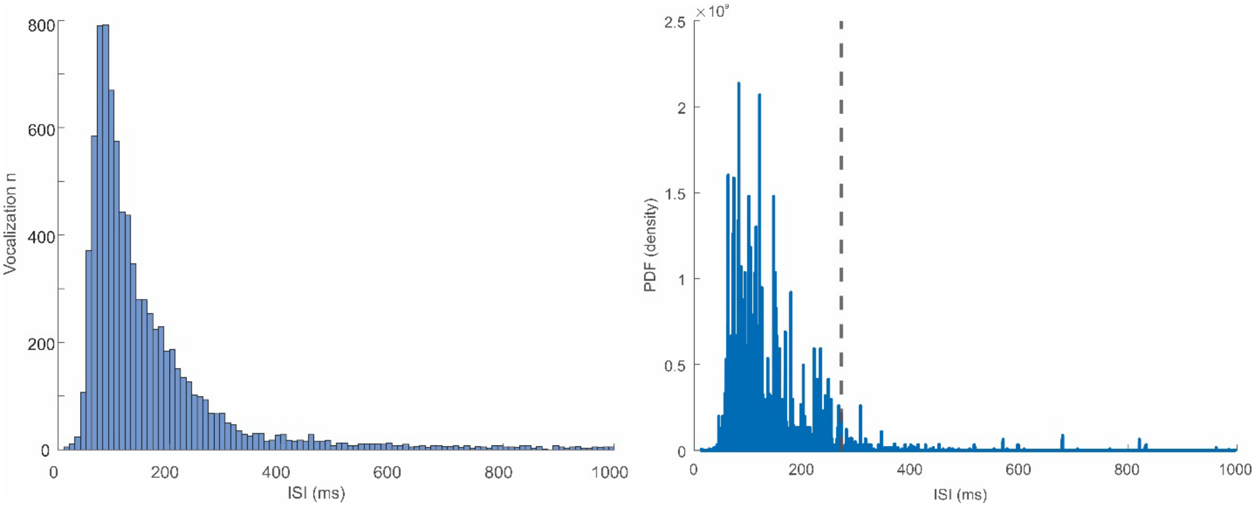


**Figure S1:** Baseline detection for call sequences. Left: histogram of inter-syllable intervals (ISIs) across all calls. Right: the probability density function (PDF) was estimated as the derivative of the cumulative distribution function (CDF). The median ISI was 117 ms, and the distribution was examined for pronounced low-density regions. The first discontinuity occurred at 271 ms, which was used as the threshold to define sequence boundaries.

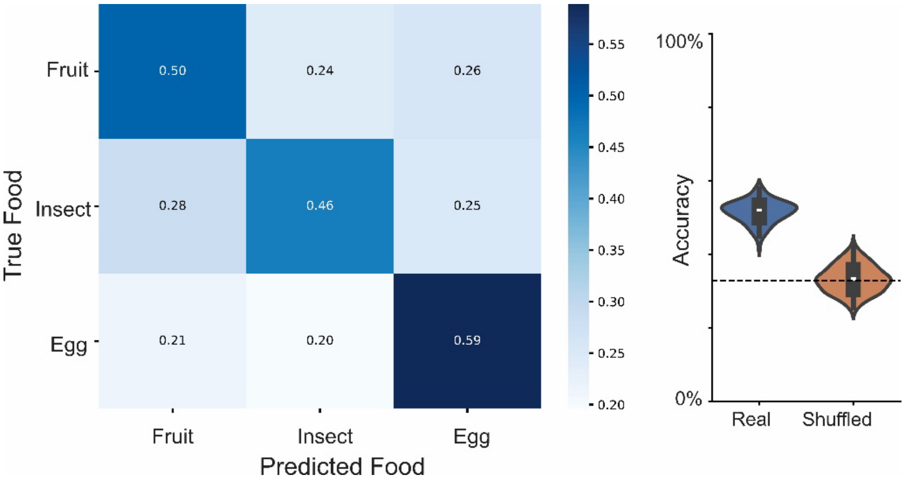


**Figure S2:** Population-level decoding of food categories for the subset of monkeys that produced calls for every category. For each iteration, 100 vocalizations per subject were sampled, and classification accuracy was computed over 100 bootstraps (mean accuracy: 52 % ± 3 %, chance: 33%).

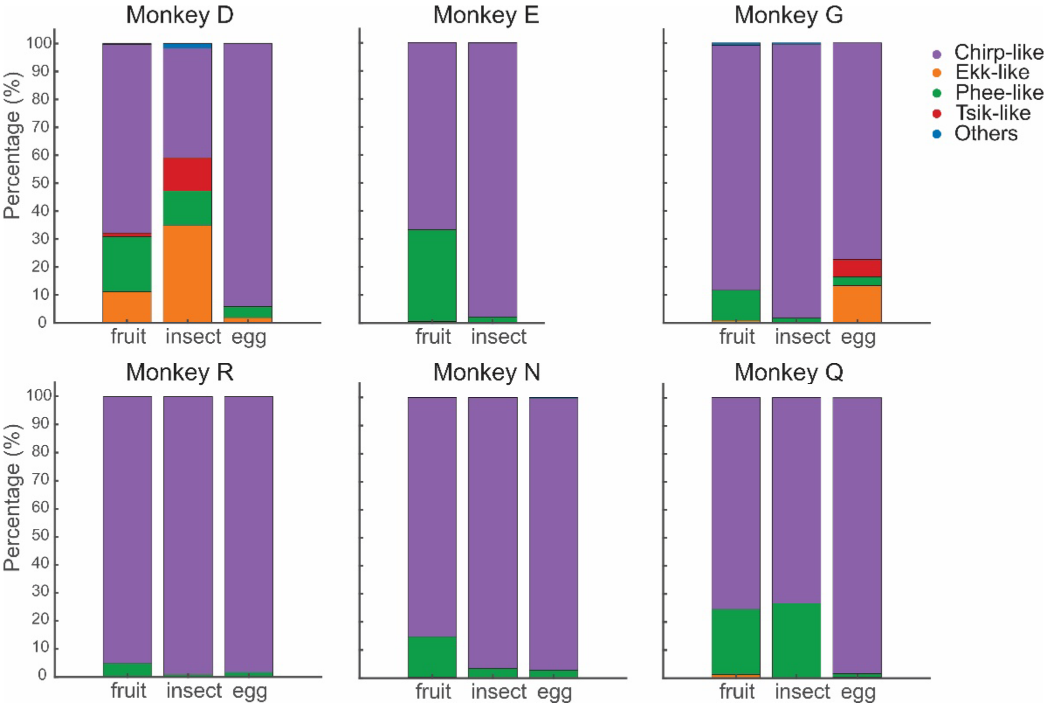


**Figure S3:** Percentages of vocalizations assigned to each cluster for each animal.

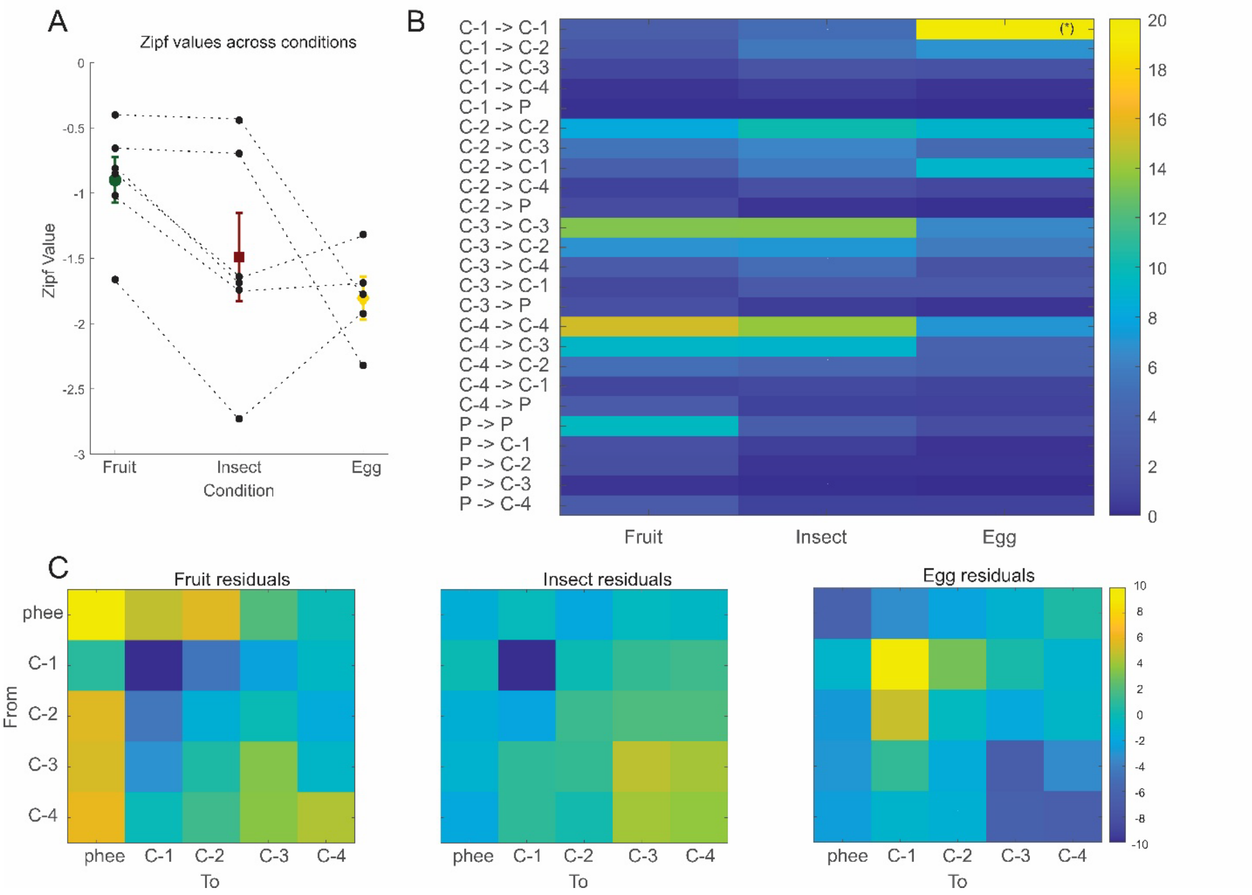


**Figure S4: Zipf values and transition dynamics across conditions. (A)** Zipf values calculated for each condition. Mean values across subjects are shown for fruit (green), insect (red), and egg (yellow) with lines indicating within-subject changes in distribution structure. **(B)** Heatmap of non-uniform transition probabilities between clusters, indicating structured sequencing rather than random combinations, and highlighting condition-specific cluster-transitions; asterisks denote values above 25%. **(C)** Chi-square residuals highlighting transitions that occur significantly more or less often than expected under independence, revealing preferred and avoided transitions in vocal sequences.

**Supplementary Tables:**

**Table S1:** Description of the features used in Fig 1D, 2D, 2E.

| Parameter | Definition |
|---|---|
| Frequency Slope | Mean peak frequency calculated across the entire vocal element. |
| Duration | Time between start and end of the vocal output. |
| Slope | Rate of change in peak frequency over time, calculated as the difference between peak frequency at the end and at the start divided by the duration of the vocalization. |
| Inter-syllable Interval (ISI) | Time difference between the end of one vocalization and the onset of the next |

**Table S2:** Food tournament results. “X” signifies when animals did not consume those items.

| Animal | First Preference | Second Preference | Third Preference |
|---|---|---|---|
| Monkey D | Locust | Banana | Egg |
| Monkey E | Grapes | Locust | X |
| Monkey G | Locust | Grapes | Egg |
| Monkey R | Locust | Egg | Banana |
| Monkey N | Locust | Grapes | Egg |
| Monkey Q | Grapes | Egg | X |

**Table S3:** Summary of Statistical Tests

| Fig 1D | Acoustic Parameter ~ Syllable Position + (Syllable Position \|SubjectNum) |
|---|---|
| Parameter | Main effect (Syllable Position) |
| z Slope Start | F(4, 3413) = 93.36, p =1.8e-75 |
| z Slope End | F(4, 3410) = 19.29, p = 1.04e−15 |
| z Peak Freq Start | F(4, 3413) = 81.75, p = 2.35e-66 |
| z Peak Freq End | F(4, 3410) = 7.18, p = 9.42e-06 |
| z Duration Start | F(4, 3413) = 9.61, p = 1.00e−07 |
| z Duration End | F(4, 3410) = 12.74, p = 2.70e−10 |
| ISI Start | F(4, 3239) = 9.06, p = 2.81e−07 |
| ISI End | F(3, 2728) = 76.16, p = 2.63e-47 |

| **Fig 2D:** | Acoustic Parameter ~ Condition + (Condition\|SubjectNum) | | |
|---|---|---|---|
| Param | Main Effect (Condition) | Condition | Post-hoc (p-value) |
| Slope | F(2, 6353)= 6.02, p=0.002 | Fruit vs Insect | 0.016 |
| | | Fruit vs Egg | 6e-04 |
| | | Insect vs Egg | ns |
| Peak Freq | F(2, 6353)= 9.13 , p=0.0001 | Fruit vs Insect | ns |
| | | Fruit vs Egg | 9.33e-04 |
| | | Insect vs Egg | ns |
| Duration | F(2, 6353)= 12.64, p= 3.3157e-06 | Fruit vs Insect | 2.24e-04 |
| | | Fruit vs Egg | 5.1635e-07 |
| | | Insect vs Egg | ns |
| ISI | F(2, 5670)= 21.03, p= 7.8856e-10 | Fruit vs Insect | 1.2378e-05 |
| | | Fruit vs Egg | 1.3789e-05 |
| | | Insect vs Egg | 0.0231 |

| Fig 2E | Acoustic parameter ~ Condition * Syllable Position + (1\|SubjectNum) | | |
|---|---|---|---|
| Parameter | Main effect (Syllable) | Main effect (Condition) | Interaction |
| z Slope Start | F(4,3403)= 124.35, p=3.39e-99 | F(2,3403)= 3.1055, p= 0.044929 | F(8,3403)=2.45, p=0.012 |
| z Slope End | F(4,3400)= 22.426, p=2.65e-18 | F(2,3400)= 0.47056, p= 0.62469 | F(8,3400)= 5.33, p = 1.14e-6 |
| z Peak freq Start | F(4,3403)= 23.19, p=9.92e-11 | F(2,3403)=31.78, p= 4.8958e-26 | F(8,3403)=2.18, p=0.026 |
| z Peak freq End | F(4,3400)= 10.767, p=1.13e-08 | F(2,3400)= 36.38, p= 2.3279e-16 | F(8,3400) = 1.75, p = 0.081 |
| z duration Start | F(4,3403)= 29.476, p=3.93e-24 | F(2,3403)= 2.467, p= 0.084987 | F(8, 3403)=5.50, p=6.22e-7 |
| z duration End | F(4,3400)= 41.397, p=6.22e-34 | F(2,3400)= 0.47557, p= 0.62157 | F(8,3400)= 9.93, p= 9.4e-14 |
| z ISI Start | F(4,3229)= 15.649, p=1.093e-12 | F(2, 3229)= 1.0899, p= 0.33638 | F(8, 3229)=6.87, p=5.3e-9 |
| z ISI End | F(3, 2720)= 24.892, p=6.94e-16 | F(2, 2720)= 5.0252, p= 0.0066311 | F(6,2720)=1.47, p=0.185 |

**Table S4:**

| **Acoustic features** | **Definition** |
|---|---|
| Peak frequency at start, center, and end | Frequency at maximum power measured at start, midpoint, and end of the vocalization. |
| Peak frequency max and min | Highest and lowest values of peak frequency within the element. |
| Peak frequency max-, min-, mean-entire | Maximum, minimum, and mean peak frequency calculated across the entire vocal element. |
| Bandwidth at start, center, and end | Difference between maximum and minimum frequencies measured at start, midpoint, and end of the vocalization. |
| Bandwidth max | Maximum bandwidth of the vocal element calculated as the difference between highest and lowest frequencies. |
| Bandwidth max-, min-, mean-entire | Maximum, minimum, and mean bandwidth calculated across the entire vocal element. |
| Bandwidth peak | Difference between peak frequency at the end and at the start of the vocal output. |
| Frequency Slope | Rate of change in peak frequency over time, calculated as the difference between peak frequency at the end and at the start divided by the duration of the vocalization. |
| Duration | Time between start and end of the vocal output. |
| Curvature | Bending of the vocalization, defined from the first and second derivate of the frequency contour calculated from peak frequency values at 5 positions of the vocalization. |
| Distance to maximum | The distance from start to the location of the maximum amplitude is measured. |
